# Repurposing protein degradation for optogenetic modulation of protein activities

**DOI:** 10.1101/680207

**Authors:** Payel Mondal, Vishnu V. Krishnamurthy, Savanna R. Sharum, Neeka Haack, Kai Zhang

**Author notes:** Corresponding author: Kai Zhang, PhD, Department of Biochemistry, School of Molecular and Cellular Biology, University of Illinois at Urbana-Champaign, 600 South Mathews Avenue, Urbana, IL 61801, USA.

## Abstract

Non-neuronal optogenetic approaches empower precise regulation of protein dynamics in live cells but often require target-specific protein engineering. To address this challenge, we developed a generalizable light modulated protein stabilization system (GLIMPSe) to control intracellular protein level independent of its functionality. We applied GLIMPSe to control two distinct classes of proteins: mitogen-activated protein kinase phosphatase 3 (MKP3), a negative regulator of the extracellular signal-regulated kinase (ERK) pathway, as well as a constitutively active form of MEK (CA MEK), a positive regulator of the same pathway. Kinetics study showed that light-induced protein stabilization could be achieved within 1 minute of blue light stimulation. GLIMPSe enables target-independent optogenetic control of protein activities and therefore minimizes the systematic variation embedded within different photoactivatable proteins. Overall, GLIMPSe promises to achieve light-mediated post-translational stabilization of a wide array of target proteins in live cells.

## Introduction

A key factor that drives the dynamic nature of signaling pathways is spatial and temporal control of protein activities. Conventional genetic and pharmacological approaches such as gene overexpression, gene knock out, RNA interference helps delineate interaction maps of signaling components. These approaches, unfortunately, lack the flexibility to resolve signaling dynamics in live cells. Light serves as an attractive tool for the regulation of signal transduction as it can be rapidly controlled with high spatiotemporal resolution^1–3^. This strategy has been widely used to control target protein activity by protein translocation^4–7^, protein caging^8–11^, sequestration^12, 13^, clustering^14, 15^, induced avidity^16–18^ or allostery^19, 20^. Successful application of non-neuronal optogenetics in multicellular organisms has provided new insights into cell fate determination during embryonic development^21–26^. Most of these systems, however, require target-specific protein engineering and often cannot be generalized to control a broader class of proteins.

A generalizable strategy for post-translational reduction of target protein level involved fusion of a degradation peptide sequence, or degron, to a protein of interest. Light-mediated uncaging of the degron results in controlled protein degradation in *Saccharomyces cerevisiae*^27^ and mammalian cells^28^. This strategy elicits in a sense a post-translational, optogenetic “knock-down” effects of a protein of interest. A comprehensive understanding of a signaling pathway would benefit from the complementary post-translational, optogenetic “knock-in” strategy, which has not been available to date.

To fill in this gap, here, we have developed a generalizable light modulated protein stabilization system (GLIMPSe) that allows for optical enhancement of protein stability. GLIMPSe consists of a degradation-rescue module which synergistically controls protein level in cells – the degradation module constantly suppresses protein level until the photo-sensitive rescue module triggers protein stabilization. Compared with previous optogenetic strategies, GLIMPSe allows for optical control of different classes of proteins and stabilizes protein level within 1 minute of blue light stimulation. To demonstrate the generalizability of GLIMPSe, we achieved optical control of functionally distinct classes of proteins including mitogen-activated protein kinase phosphatase 3 (MKP3) and a constitutively active MEK (CA MEK). Thus, GLIMPSe enables bidirectional control of the extracellular signal-regulated kinase (ERK) signaling pathway, which regulates crucial cell functions such as proliferation, differentiation, migration, and apoptosis. We expect that GLIMPSe would add a powerful capacity to the current optogenetics toolbox and promise to lower the technical barrier for achieving optical control of protein activities.

## Results

### Reduce the intracellular protein level by a degron from the Deadend protein

The Deadend (Dnd) protein regulates germline development in vertebrate. In mice, cells lacking Dnd fail to support germline development^29^. We have recently discovered a 25 amino-acid degradation sequence within *Xenopus laevis* Dnd protein that mediates stage-dependent degradation of Dnd during Xenopus embryonic development. To determine whether this degradation sequence (here referred to as degron or deg) can be used as a general moiety to mediate degradation of intracellular proteins, we fused tandem arrays of degrons to the C-terminus of firefly luciferase (FLuc) protein (FLuc-tevS-Ndeg, N=1, 2, and 3). A tobacco etch virus (TEV) protease recognition site (tevS) was sandwiched between FLuc and degron to render TEV-mediated rescue of protein degradation (**Fig. 1a**). HEK293T cells were cotransfected with FLuc-tevS-Ndeg (N=1, 2, 3) and a constitutively expressed Renilla luciferase (RLuc, loading control), recovered overnight and harvested for a dual-luciferase analysis. Degrons significantly reduced the luciferase signals in a dose-dependent way, with N=3 showing the optimal contrast compared to the no-degron control (**Fig. 1b**). Thus, we used three tandem degrons (denoted as deg) in all other constructs. When HEK293T cells were cotransfected with plasmids encoding TEV protease, a 5-fold increase of FLuc signal was observed compared with the no-TEV control (**Fig. 1b**).

**Figure 1.**
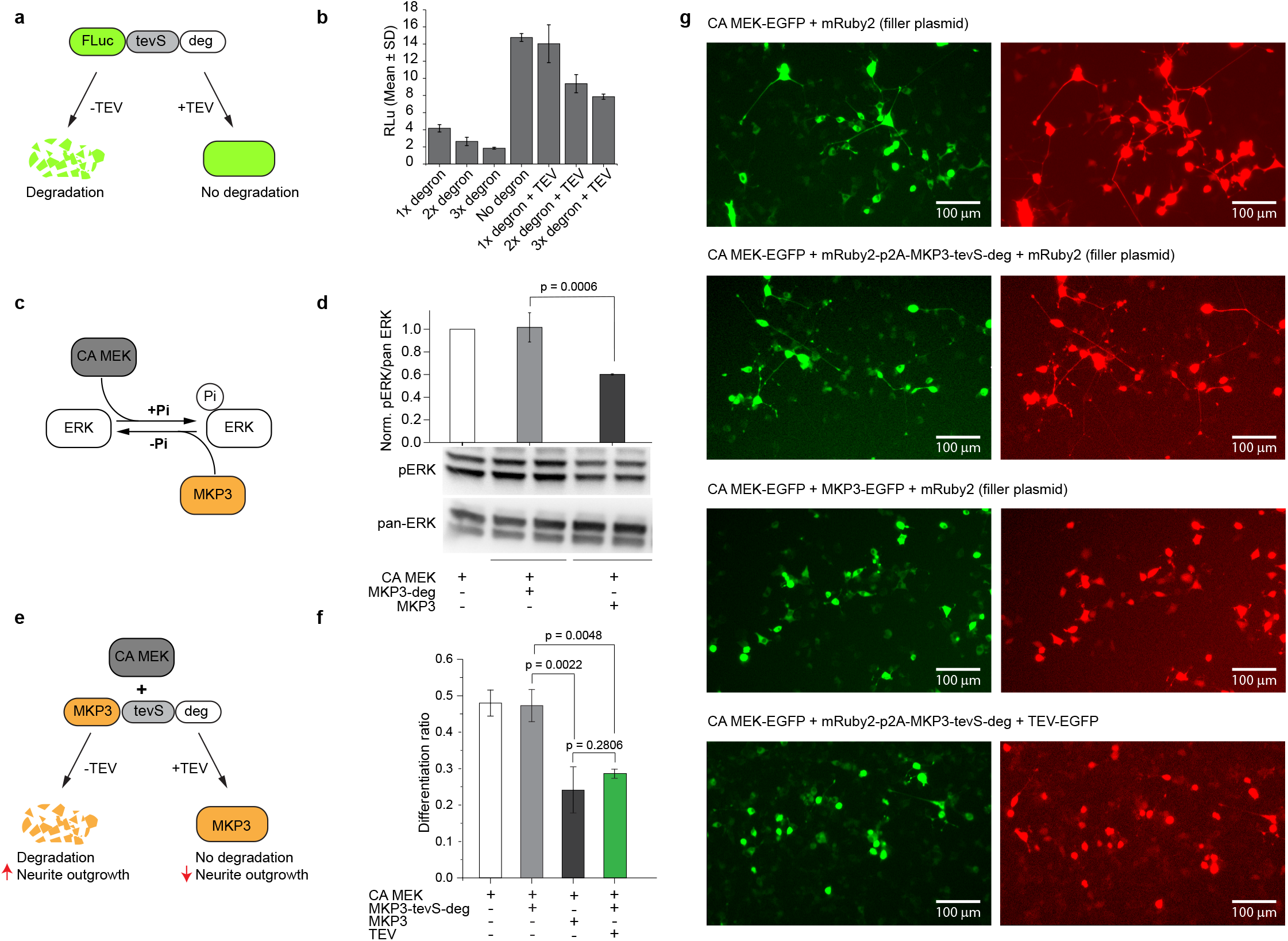
Control intracellular protein level with degron. (a) Schematic for TEV protease-mediated rescue of firefly luciferase (FLuc) protein degradation. (b) Degradation of FLuc-tevS-deg and TEV-mediated protein stabilization demonstrated by a dual-luciferase assay in HEK293T. (c) Schematic for bidirectional regulation of ERK by CA MEK and MKP3, respectively. (d) Western blot analysis of pERK/pan-ERK level in PC12 cells under different conditions. The pERK/pan-ERK intensity ratio was normalized to that of CA MEK overexpression (left-most lane). (e) Schematic for regulating MKP3 stability with degron and TEV protease. (f) Quantification of PC12 cell differentiation ratio under different conditions. Values represent Mean ± SD of three biological replicates (n = 3). More than 200 cells were counted per replicate. (g) Representative images of PC12 cells under different conditions. The total amount of plasmids was kept the same between each condition by including an mRuby2 plasmid as a filler. Scale bar 100 μm.

### TEV-mediated rescue of MKP3 degradation inhibits the ERK pathway

To determine if this degron system can be used to control the intracellular signaling outcome, we proceeded to determine the efficiency of degron with mitogen-activated protein kinase phosphatase 3 (MKP3). MKP3 is a cytoplasmic phosphatase that specifically dephosphorylates the phospho-ERK1/2 (pERK1/2)^30^. To assess if degron could modulate MKP3 activity, we used a neurite outgrowth assay in the rat PC12 pheochromocytoma cell line.

To visualize degron fused MKP3 transfected PC12 cells, we fused an mRuby2 fluorescent protein into the same vector with a p2A peptide (mRuby2-p2A-MKP3-tevS-deg). p2A is a consensus motif that results in efficient ribosomal skipping between two cistrons in mammalian cells^31^. Insertion of the p2A peptide allows only MKP3 to undergo a constitutive degradation, leaving mRuby2 intact for cell visualization.

To better show the effect of MKP3, we first hyperactivated the ERK pathway by transfecting PC12 cells with a constitutively active dual specificity mitogen-activated protein kinase (CA MEK)^32^ (**Fig. 1c**). CA MEK enhanced the pERK level in PC12 cells, as evidenced by Western blot analysis. Upon fusion with degron, MKP3 underwent a baseline degradation, and the ERK pathway remained hyperactivated compared with the no-degron control (**Fig. 1d, bar 2 and 3**). Single-cell analysis of PC12 cell differentiation showed a consistent result, where MKP3-deg did not reduce CA MEK-mediated neurite outgrowth (**Fig. 1e, 1f bar 1 and 2**) in contrast with MKP3 (**Fig. 1f, bar 3**). When TEV was co-transfected with MKP3-tevS-deg and CA MEK, MKP3 degradation was successfully rescued, which resulted in a reduced PC12 cell differentiation ratio comparable to that of MKP3 (**Fig. 1f, bar 4**). Representative fluorescence images under various transfection conditions were shown in **Fig. 1g**.

### Use GLIMPSe to inhibit the ERK pathway via light-induced stabilization of MKP3

To develop an optogenetic system for light-induced stabilization of MKP3, we adopted a two-module strategy. First, MKP3 is constantly degraded by the degradation module. Second, an optogenetic strategy is used to supply the functional TEV protease, offering a photosensitive degradation-rescue module. We chose to modify the recently developed light-inducible nuclear export system (LEXY) as the degradation-rescue module^9^. LEXY consists of a C-terminal caged nuclear export signal (NES) and an N-terminal nuclear localization signal (NLS) to accumulate the fusion protein into the nucleus in the dark. Blue light uncages NES and mediates active nuclear export through the nuclear pores.

We fused TEV into the LEXY system (TEV-LEXY) so that the TEV protease would be sequestered into the nucleus in the dark and exported into the cytoplasm in response to blue light (**Fig. 2a**). However, PC12 cells co-transfected with MKP3-tevS-deg and TEV-LEXY showed no significant difference in their differentiation ratio (0.23 vs. 0.24) (**Fig. 2b, bar 1 and 2**). Reduced differentiation ratio in the absence of light may arise from leaking of TEV-LEXY into the cytoplasm, which cleaves tevS and stabilizes MKP3. To address this issue, we caged the tevS with a recently developed evolved LOV (eLOV) protein from *Avena sativa*^33^ (MKP3-eLOVtevS-deg) (**Fig. 2a**). Addition of the eLOV successfully protected tevS cleavage (therefore MKP3 degradation) and resulted in an almost full recovery of the PC12 cell differentiation ratio in the dark (**Fig. 2c, bar 2**). In response to blue light stimulation, uncaging of eLOV exposed the tevS so that it can be cleaved by TEV protease, resulting in MKP3 recovery and a reduced PC12 cell differentiation ratio (**Fig. 2c, bar 1**). Light inactivation of ERK pathway in PC12 cells was also validated by 1.9-fold reduction of pERK level (**Fig. 2d, e**) assayed by Western blot.

**Figure 2.**
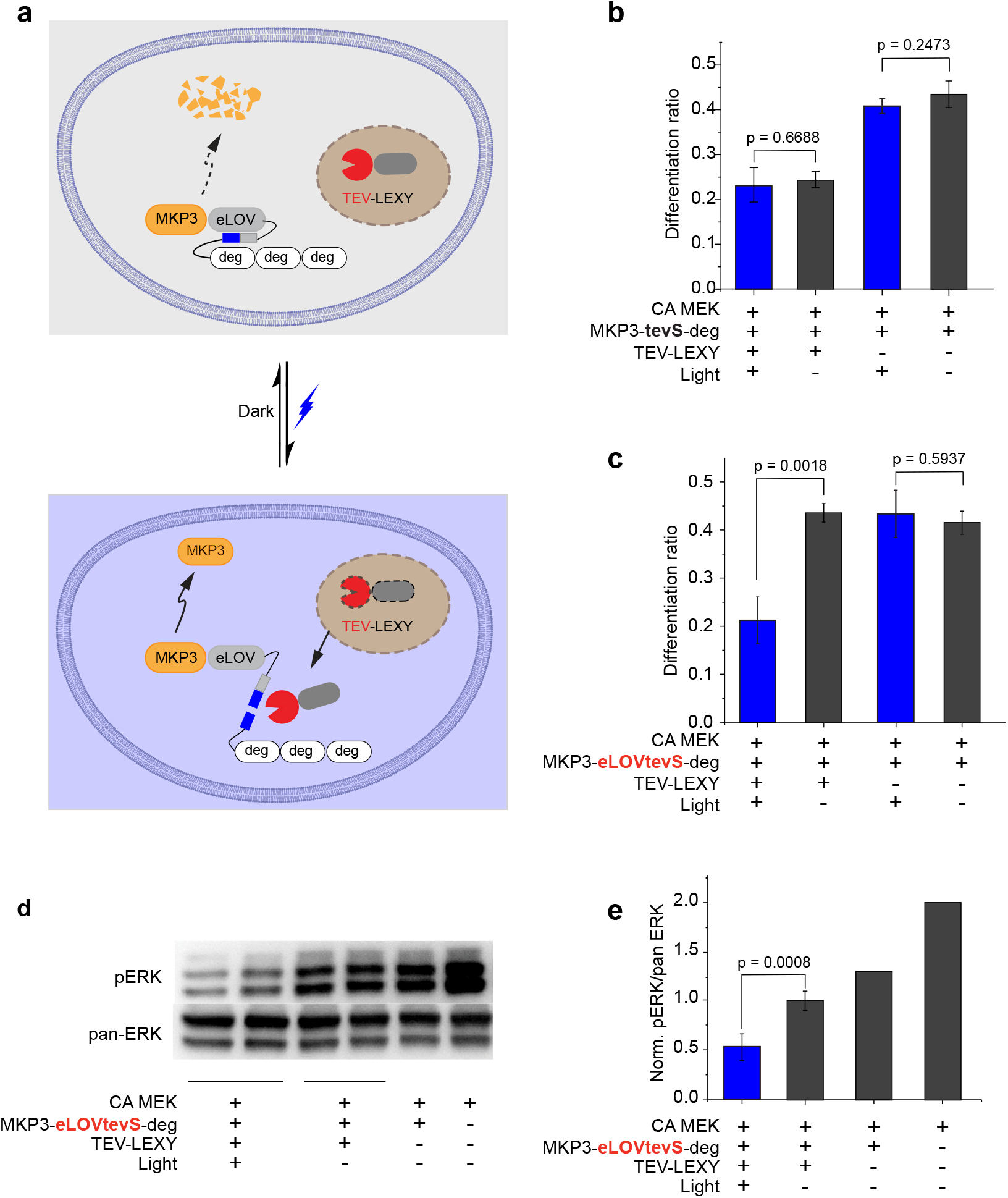
Generation of GLIMPSe for light-induced MKP3 stabilization. (a) Schematic for optogenetic control of MKP3 stability with GLIMPSe. (b) Quantification of PC12 cell differentiation ratio under different conditions. Cells were illuminated with 500 μW/cm^2^ blue light or kept in the dark for 45 h prior to imaging. Values represent Mean ± SD of three biological replicates (n = 3) with more than 200 cells counted per replicate. (c) Quantification of differentiation ratio for PC12 cells expressing GLIMPSe-MKP3 optogenetic system. Differentiation ratio = # of differentiated cells / # of transfected cells. The bar graph is presented with Mean ± SD averaged over three independent experiments (n=3). (d) Western blot analysis of PC12 cells expressing CA MEK, MKP3-eLOVtevS-degron and TEV-LEXY. (e) Normalized intensity of pERK/pan-ERK in PC12 cells after light-induced MKP3 stabilization. Band intensities are normalized to the average intensity of lane 3 and 4. The bar graph is presented with Mean ± SD averaged over four replicates (n=4).

### GLIMPSe functions within 1 minute of blue light stimulation

To determine the kinetics of optical control of MKP3 stabilization, we varied the duration of blue light stimulation. We first determined the response time of TEV-LEXY with optical microscopy and found cytosolic export of TEV-LEXY occurs within 5 min of blue light exposure (**Fig. 3a**). Thus, we designed two series of light duration ranging from 1 min to 2 h to resolve fine kinetics (**Fig. 3b, top 2 panels**) and from 0.5 to 24 h to examine long-term effect (**Fig. 3b, bottom 2 panels**). Because the cleaved protein loses three degrons, its molecular weight reduces approximately 10 kDa compared with the uncleaved form (e.g., the size of HA-MKP3-eLOVtevS-deg is 72 kDa, whereas the size of HA-MKP3-eLOV is 62 kDa). The protein stabilization efficiency was therefore calculated as

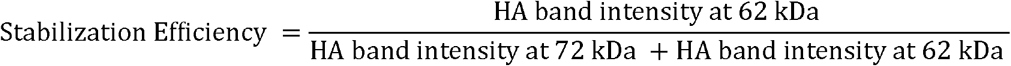

**Fig. 3.**
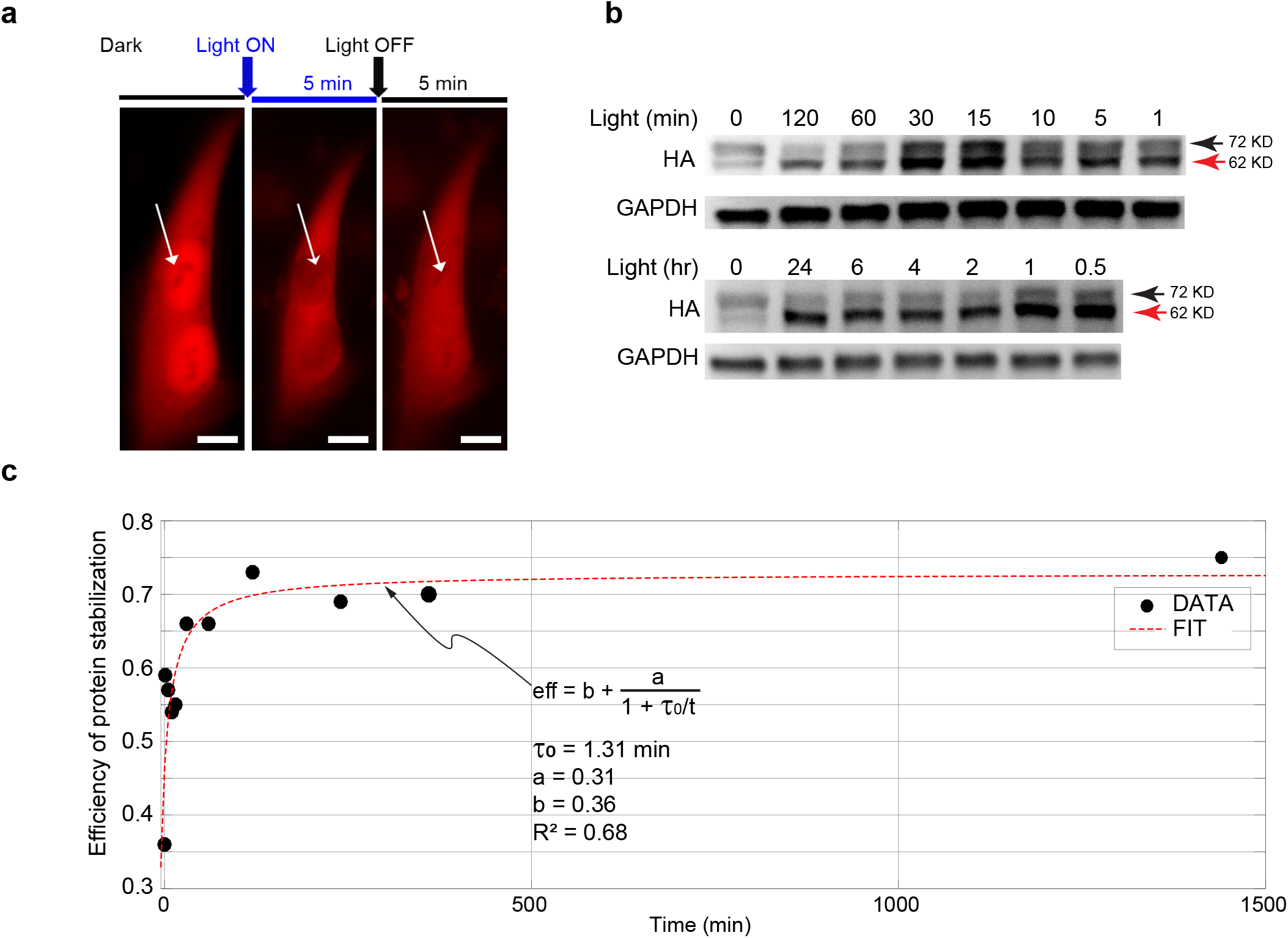
Kinetics of blue light-induced protein stabilization. (a) Kinetics of TEV-LEXY nuclear-cytoplasmic shuttling upon 5 min of blue light stimulation in HEK293T cells. Scale bar: 10 μm. (b) Kinetics of light-induced protein stabilization in PC12 cells with 0.5 mW/cm^2^ blue light stimulation probed by a gel-shift assay. Cells kept in the dark showed minimum size shift from 72 kDa to 62 kDa, indicating that most of the HA-MKP3-eLOVtevS fusion protein was still fused to degron. Within 1 min of blue light stimulation, a strong size shift was observed, which was maintained up to 24 h after light stimulation. (c) The efficiency of protein stabilization in response to blue light stimulation. Protein stabilization efficiency in each time point was calculated by dividing the band intensity at 62 kDa with the sum of 72-kDa and 62 kDa band intensity in the corresponding lane. The efficiency vs. time curve was then fit in MATLAB.

We observed significant band shift from 72 kDa to 62 kDa within only 1 min of blue light exposure and a strong size shift was maintained with increasing time for blue light stimulation up to 24 h (**Fig. 3b**). The efficiency vs. time curve was then fit in Matlab by the following equation (**Fig. 3c**)

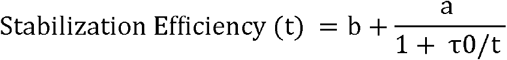

where “b” represents basal activity, “a” represents the maximum enhancement of cleavage activity from the basal level, and τ0 represents the time to reach half the maximum of a. The fit result shows that optical control of protein stabilization occurs on the order of 1 min of blue light stimulation.

### Use GLIMPSe to activate the ERK pathway by light-induced CA MEK stabilization

To demonstrate the generalizability of GLIMPSe, we set out to control a different class of protein from MKP3. We chose a constitutively active dual specificity mitogen-activated protein kinase (CA MEK) that activates the ERK pathway. Before the construction of GLIMPSe-CA MEK, we verified that CA MEK could be degraded by degron and its degradation could be rescued by cotransfection of TEV protease, which was assayed by PC12 cell differentiation and Western blot (**Fig. 4**). PC12 cells transfected with CA MEK-tevS-deg showed a significant reduction of differentiation ratio compared to cells transfected with CA MEK (**Fig. 4a, b**), which was consistent with the 2.17-fold reduction of the pERK activity (**Fig. 4c**). Co-transfection of TEV protease with CA MEK-tevS-deg successfully rescued CA MEK degradation and resulted in 2.67-fold increased PC12 differentiation ratio (**Fig. 4d-e**) as well as the 1.44-fold elevation of pERK level compared to cells without TEV-EGFP (**Fig. 4f**). To construct GLIMPSe-CA MEK (**Fig. 5a**), we replaced MKP3 in GLIMPSe-MKP3 with CA MEK. PC12 cells transfected with GLIMPSe-CA MEK and exposed to 45-h blue light stimulation showed a 2.5-fold increase in their differentiation ratio (**Fig. 5b, bar 3 and 4**) and 2-fold elevation of pERK level compared to cells kept in the dark (**Fig. 5c, d**). Negative controls without TEV-LEXY showed base-level differentiation ratio (**Fig. 5b, bar 1 and 2**) as well as pERK level (**Fig. 5d, bar 1 and 2**).

**Figure 4.**
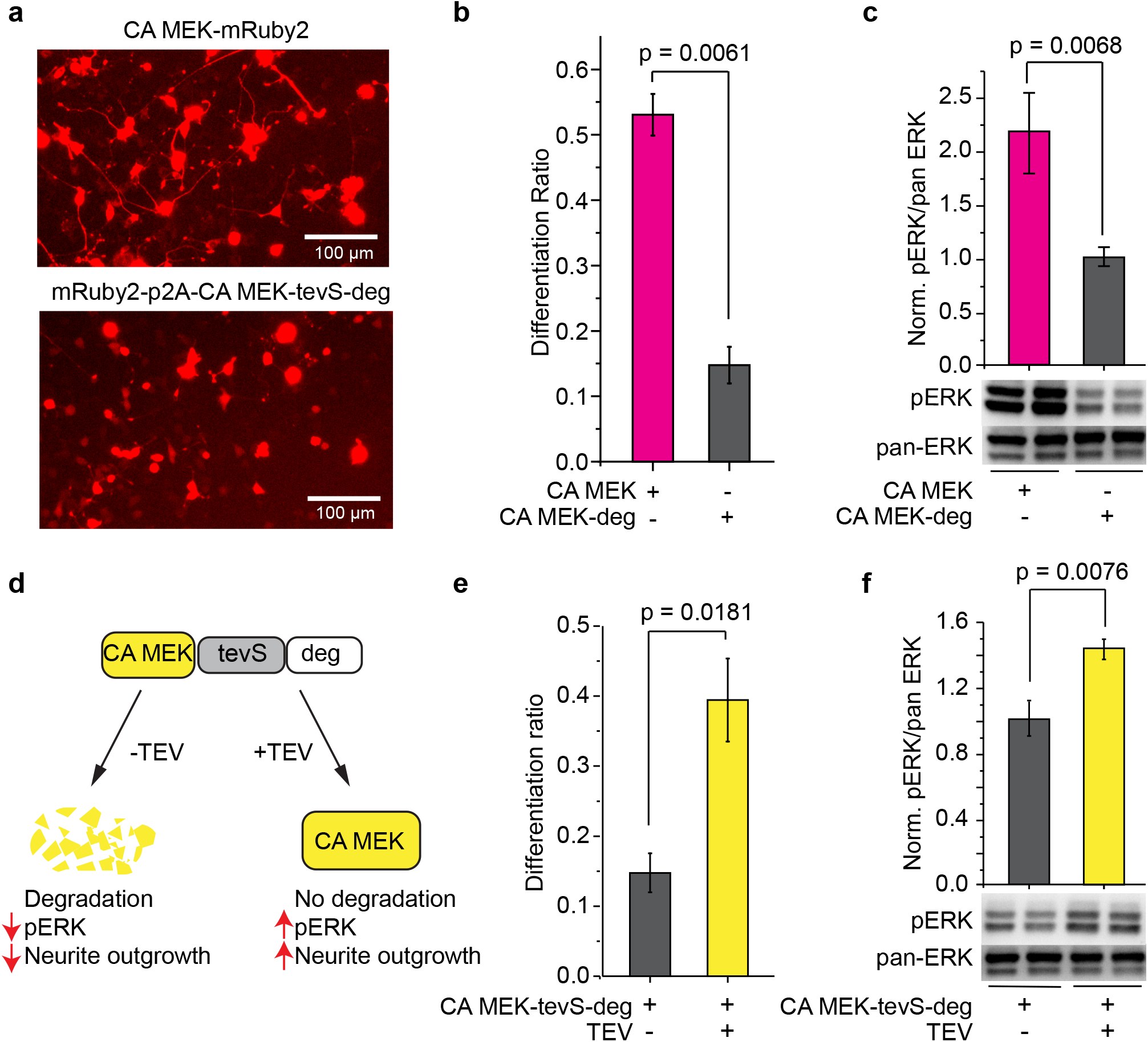
Control the intracellular level of CA MEK with degron and TEV. (a) Representative images of PC12 cells transfected with CA MEK-mRuby2 (top) or mRuby2-p2A-CA MEK-tevS-degron (bottom). Scale bar: 100 μm. (b) Cells transfected with degron-fused CA MEK showed significantly less neurite outgrowth than those transfected CA MEK. (c) Western blot analysis of pERK/pan-ERK level in PC12 cells transfected with CA MEK or CA MEK-deg. Values at each condition were normalized to the intensity of CA MEK-degron. The bar graph is presented with Mean ± SD averaged over three independent sets of experiments (n=3). (d) Schematic for controlling CA MEK protein stability with C-terminal fused degron and TEV protease. (e) Quantification of PC12 cell differentiation ratio under different conditions. PC12 cells transfected with mRuby2-p2A-CAMEK-tevS-degron showed significantly less differentiation, which was rescued by co-transfection of TEV-EGFP. (f) Western blot analysis for TEV-mediated rescue of CA MEK stability and elevation of pERK level in PC12 cells. pERK/pan-ERK intensity was normalized to the intensity of CA MEK-tevS-degron in the absence of TEV protease. The bar graph is presented with Mean ± SD averaged over three replicates (n=3).

**Figure 5.**
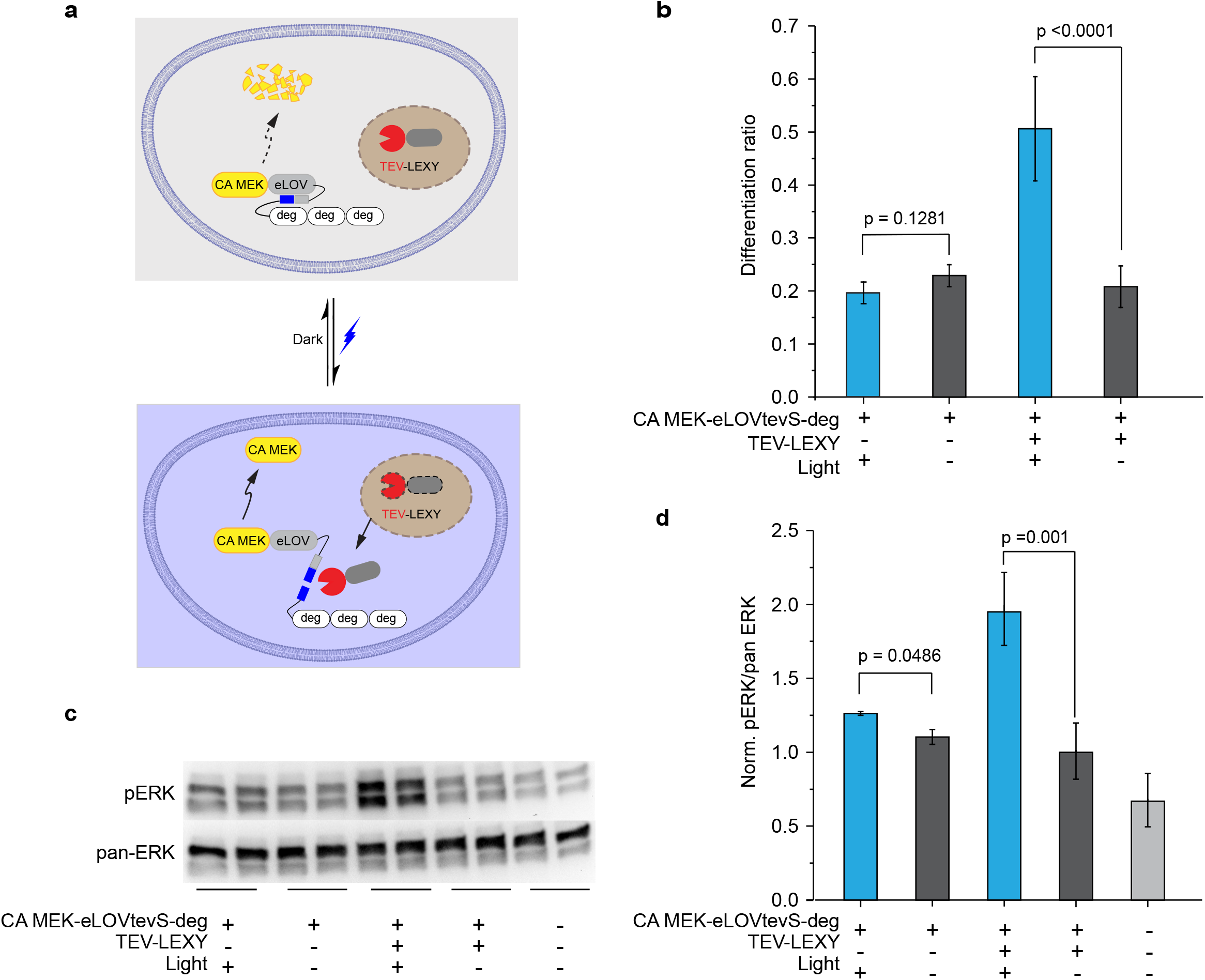
Generation of GLIMPSe for light-induced CA MEK stabilization. (a) Schematic for optogenetic control of CA MEK stability with GLIMPSe. (b) Quantification of differentiation ratio for PC12 cells expressing GLIMPSe-CA MEK optogenetic system. Cells were illuminated with 500 μW/cm^2^ blue light or kept in the dark for 45 h prior to imaging. The bar graph is presented with Mean ± SD averaged over three biological replicates (n = 3) with more than 200 cells counted per replicate. (c) Western blot analysis of PC12 cells expressing GLIPMSe-CA MEK under dark and light. (d) Normalized intensity of pERK/pan-ERK level in PC12 cells transfected with GLIMPSe-CA MEK. Band intensities are normalized to the average intensity of lane 7 and 8. The bar graph is presented with Mean ± SD averaged over four biological replicates (n = 4).

## Discussion

Development of a generalizable optogenetic platform would significantly lower the technical barrier for the utilization of non-neuronal optogenetics. For example, a recent computational framework has been developed to facilitate the design of split proteins for optogenetic control of protein activities^34^. In this study, we have developed a generalizable light modulated protein stabilization system (GLIMPSe) that allows for optical control of protein stability in live cells. By integrating a degron derived from the *Xenopus laevis* Dnd protein into two optogenetic modules (LEXY and eLOV), we can rescue degron-mediated protein degradation with a short pulse of blue light (1 min) at an amenable power dosage (0.5 mW/cm^2^). An attractive feature of the GLIMPSe system is that each regulatory module (degron, eLOV, and TEV-LEXY) does not interrupt the functionality of protein of interest and therefore can be utilized to control the intracellular level of a wide range of target proteins. To demonstrate this point, we used GLIMPSe to control two different classes of proteins: a phosphatase (MKP3) that acts as a negative regulator of ERK pathway and a kinase (CA MEK), an activator of the same pathway.

Taking advantages of the high spatiotemporal resolution as well as its capacity to delineate signaling sub-circuits, optogenetic technology promises to offer new insights into signal transduction in live cells. It has been increasingly realized, however, that quantitative analysis of signaling outcomes suffers from systematic variations embedded within different photoactivatable proteins such as binding affinity, dynamic range, as well as the expression level. GLIMPSe offers a platform that allows for target-independent optogenetic control of protein activities and therefore minimizes the systematic variation. More importantly, each functional unit (e.g., eLOV and LEXY) in GLIMPSe is modular, and their functionality is tunable. Thus, GLIMPSe would retain a high upgrading potential and help promote the use of nonneuronal optogenetics by reducing the cost and time for target-specific protein engineering. Although GLIMPSe requires blue light that has a limited penetration depth in biological tissues, we believe that recent development of wireless optogenetics based on emerging nanomaterials such as upconversion nanoparticles would address this challenge^35–37^.

## Materials and Method

### Reagents

Phusion DNA polymerase master mix (M0530L) was purchased from NEB. In-Fusion HD Cloning Plus kit was from Takara (638909). DreamTaq PCR Master Mix (2×) (K1081), Turbofect (R0532) and Pierce Protease and Phosphatase Inhibitor Mini Tablets (A32959), F12K (21127-022) medium, and horse serum (26050088) were from Thermo Fisher Scientific. Fetal bovine serum (F1051), RIPA Lysis Buffer, 10× (20-188) was from Millipore Sigma. PBS (21-040-CV), DMEM (10-017-CV), Penicillin-streptomycin solution (30-002-CI), Trypsin EDTA (0.25% Trypsin, 0.1% EDTA) 1× (25-053-CI) was from Corning. Precast protein gels (456-1024) and ECL substrate (170-5060) were from Bio-Rad. Antibodies used in this work are Phospho-p44/42 MAPK (Erk1/2) (Thr202/Tyr204) (Cell Signaling Technology, 9101S, 1:1000), p44/42 MAPK (Erk1/2) (Cell Signaling Technology, #9102S, 1:1000), HA-Tag (C29F4) (Cell Signaling Technology, #3724S, 1:1000), GAPDH (14C10) (Cell Signaling Technology, #2118S, 1:1000) and Anti-rabbit IgG, HRP-linked Antibody (Cell Signaling Technology, 7074S, 1:2000).

### Plasmid construction

Constitutive active MEK (S218D, S222D) was constructed by site-directed mutagenesis. Evolved LOV (eLOV) was amplified from a synthetic gBlock from IDT based on sequence from the previous work^24^. The degron sequence (GRLYEFRLMMTFSGLNRGFAYARYS) was a gift from Dr. Jing Yang at the University of Illinois Urbana-Champaign. LEXY was amplified from NLS-mCherry-LEXY (Addgene, catalog #72655). Full sequence of GLIMPSe was shown in **Fig. S3**.

### Cell culture and transfection

HEK293T cells were cultured in DMEM supplemented with 10% fetal bovine serum and penicillin-streptomycin. PC12 cells were cultured in F12K medium supplemented with 15% horse serum and 2.5% FBS. All cell cultures were maintained in a standard incubator at 37 °C with 5% CO_2_. HEK293T and PC12 cells were seeded in 12 well plates, and cells were transfected once they reached 60% −80% confluency. Transfection was performed using Turbofect transfection reagent following the vendor’s instruction. For light-induced stabilization of MKP3 experiments, 100 ng CA MEK-EGFP, 200 ng EGFP-p2A-MKP3-eLOVtevS-deg and 700 ng of NLS-mCherry-TEV-LEXY plasmid were cotransfected in PC12 cells. For light-induced CA MEK stabilization experiments 50 ng of EGFP-p2A-CA MEK-eLOVtevS-deg and 950 ng of NLS-mCherry-TEV-LEXY plasmid was cotransfected in PC12 cells. After 3 hours of transfection, transfection medium was replaced with growth medium (F12K + 15% horse serum +2.5% FBS). After overnight recovery, the cell culture was exchanged to a low-serum medium (1.5% horse serum + 0.025% FBS) for another 24 hours to reduce the base-level ERK activity.

### Long-term light stimulation for PC12 cells neurite outgrowth assay

After 3 hours of transfection, cells were switched to growth media (F12K + 15% horse serum +2.5% FBS) and exposed to 0.5 mW/cm^2^ blue light stimulation. Both the LED device and the cell culture plate were placed into a CO_2_ incubator. After overnight recovery in complete growth medium, cell culture was exchanged to low-serum starvation medium (F12K + 0.15% horse serum + 0.025% FBS), followed by another 24 hours of continuous blue light stimulation. Cells that were transfected but kept in the dark were used as a negative control. Neurite outgrowth was quantified by the end of light stimulation.

### Western Blot

After transfection, recovery, and starvation, cells were harvested and lysed with a mixture of RIPA buffer and protease/phosphatase inhibitor cocktail. Lysates were centrifuged, and supernatants were mixed with NuPAGE™ LDS Sample Buffer and β-Mercaptoethanol. Samples were subjected to SDS-PAGE on a precast gel, followed by overnight transfer to PVDF membrane. Two different blots were run using the same cell lysate to probe the phospho-ERK and pan ERK, respectively.

### Kinetic analysis of GLIMPSe-mediated protein stabilization

PC12 cells were plated and transfected in a 12-well tissue culture plate with 100 ng EGFP-p2A-HA-MKP3-eLOVtevS-deg and 50 ng of NLS-mCherry-TEV-LEXY plasmid. After 3 hours of transfection, cells were recovered overnight in growth medium (F12K + 15% horse serum +2.5% FBS). Next day, cells were exposed to 0.5 mW/cm^2^ blue light stimulation for different timespan ranging from 1 min to 24 hours. After light exposure, cells were harvested for Western blot analysis with HA tag and GAPDH primary antibody.

### Construction of a programmable LED device

For both long-term and short-term light illumination the LED array was constructed by assembling a 6-by-4 blue LED array with 24 blue LEDs (B4304H96, Linrose Electronics) on a breadboard. LED intensity can be continuously tuned through a tunable voltage and a current-limiting resistor. The breadboard was hosted in an aluminum box, and a light diffuser film was positioned above the LED array to make the light intensity homogeneous in the defined area. The light intensity at the cell culture plate was measured by a power meter (PM100D, S121C, Thorlabs).

### Statistical analysis

The p-values were determined by performing two-tailed, unpaired t-test using the GraphPad Prism software.

## Supporting information

Supplemental Images

## Acknowledgments

K.Z. thanks the support from the School of Molecular and Cellular Biology at UIUC. We also thank Dr. Tobias Meyer from Stanford University for the gift of PC12 NS1 cells, Dr. Lin-Feng Chen (UIUC) for the gift of HEK293T cells. We thank the technical support from Ranajay Mandal from Purdue University in the construction of the LED light box.

## Author Contributions

P.M. and K.Z. conceived the experimental design. P.M., V.V.K., S.R.S, and N.H. conducted the experiments. P.M. and N.H. analyzed the data. P.M. and K.Z. prepared the manuscript and figures.

## Competing interests

The authors declare no competing interests.

## Data availability

The data that support the findings of this study are available from the corresponding author upon reasonable request

