## Supplemental Images for "Repurposing protein degradation for optogenetic modulation of protein activities"

**Supplementary Figure 1**

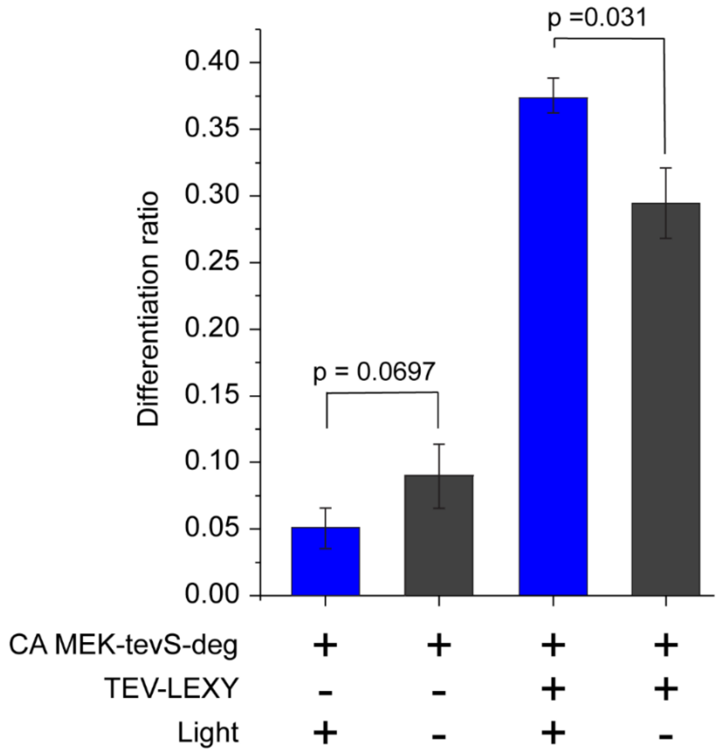


**Figure S1. Absence of eLOV shows increased stabilization of CA MEK and increased PC12 differentiation ratio in dark.** Differentiation ratio calculated for PC12 cells transfected with mRuby2-p2A-CA MEK-tevS-3X degron and NLS-mCherry-TEV-LEXY. Cells were illuminated with 500 μW/cm$^2$ blue light or kept in the dark for 45 h prior to imaging. Values represent the mean ± SD of three biological replicates (n = 3) with >100 cells counted per replicate. Differentiation ratio= (# of transfected + differentiated cells) / # of transfected cells

**Supplementary Figure 2**

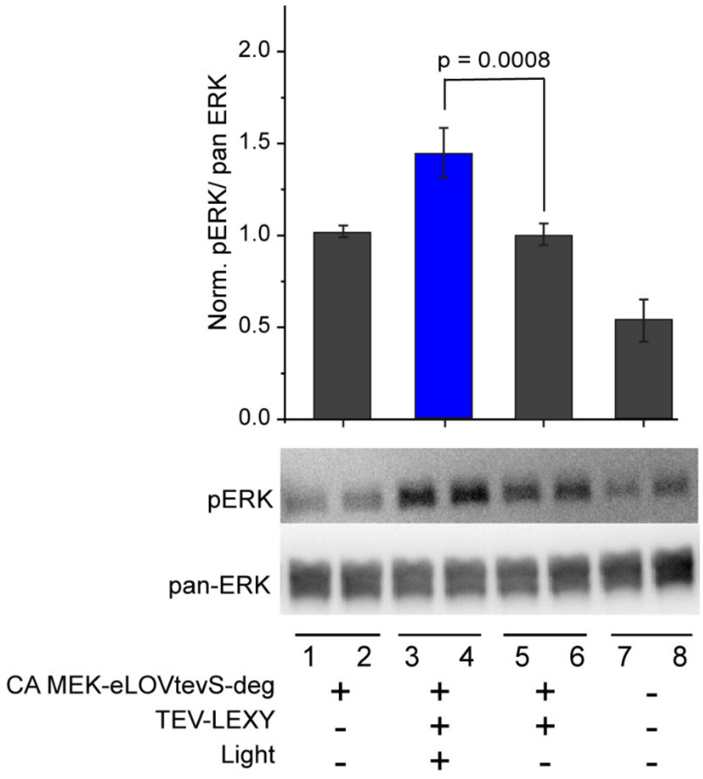


**Figure S2. Light-induced activation of GLIMPSe-CA MEK elevates the pERK level in HEK293T cells.** Western blot analysis of HEK293T cells expressing GLIMPSe-CA MEK under different conditions. Cells were kept in dark or 500 μW/cm$^2$ blue light for 24 h followed by cell lysis. Phospho-ERK/pan ERK band intensities were normalized to the average intensity of lane 5 and 6. The bar graph is presented with Mean ± SD averaged over four biological replicates (n = 4).

**Supplementary Figure 3**

```
  1 TCTAGGGCTACTACACTTGAACGTATTGAGAAGAGTTTTGTCATTACTGACCCAAGATTG 60
  1  S  R  A  T  T  L  E  R  I  E  K  S  F  V  I  T  D  P  R  L  20

 61 CCAGATAATCCCATTATATTCGTTTCCGATAGTTTCTTGCAGTTGACAGAATATAGCCGT 120
 21  P  D  N  P  I  I  F  V  S  D  S  F  L  Q  L  T  E  Y  S  R  40

121 GAAGAAATTTTGGGAAGAAACTGCAGGTTTCTACAAGGTCCTGAAACTGATCGCGCGACA 180
 41  E  E  I  L  G  R  N  C  R  F  L  Q  G  P  E  T  D  R  A  T  60

181 GTGAGAAAAATTAGAGATGCCATAGATAACCAAACAGAGGTCACTGTTCAGCTGATTAAT 240
 61  V  R  K  I  R  D  A  I  D  N  Q  T  E  V  T  V  Q  L  I  N  80

241 TATACAAAGAGTGGTAAAAAGTTCTGGAACCTCTTTCACTTGCAGCCTATGCGAGATCAG 300
 81  Y  T  K  S  G  K  K  F  W  N  L  F  H  L  Q  P  M  R  D  Q  100

301 AAGGGAGATGTCCAGTACTTTATTGGGGTTCAGTTGGATGGAACTGAGAGGGTCCGAGAT 360
101  K  G  D  V  Q  Y  F  I  G  V  Q  L  D  G  T  E  R  V  R  D  120

361 GCTGCCGAGAGAGAGGCTGTCATGCTGGTTAAGAAAACTGCAGAAGAAATTGATGAGGCG 420
121  A  A  E  R  E  A  V  M  L  V  K  K  T  A  E  E  I  D  E  A  140

421 GCAAAAgagaacctgtacttccagatgGGTGGAGGCTCTGGTAGACTCTATGAATTTAGG 480
141  A  K  E  N  L  Y  F  Q  M  G  G  G  S  G  R  L  Y  E  F  R  160

481 TTGATGATGACCTTCTCCGGGCTCAATCGCGGTTTTGCATACGCACGGTACAGTGGATCC 540
161  L  M  M  T  F  S  G  L  N  R  G  F  A  Y  A  R  Y  S  G  S  180

541 GCTAGCGGTAGACTCTATGAGTTTAGACTGATGATGACATTCTCTGGACTTAACAGAGGG 600
181  A  S  G  R  L  Y  E  F  R  L  M  M  T  F  S  G  L  N  R  G  200

601 TTCGCCTATGCCCGATATTCTGGATCCGGTAGGCTTTATGAGTTTCGCCTGATGATGACA 660
201  F  A  Y  A  R  Y  S  G  S  G  R  L  Y  E  F  R  L  M  M  T  220

661 TTTTCCGGGTTGAACAGGGGCTTCGCTTATGCTCGCTACTCAtag 705
221  F  S  G  L  N  R  G  F  A  Y  A  R  Y  S  *  235
```

Evolved LOV (eLOV) | TEV recognition site (tevS)
Degron | Degron | Degron

**Figure S3. DNA and amino acid sequences of GLIMPSe.** The domain of evolved LOV domain, tevS, and three codon-optimized degron are marked in distinct colors. Protein of target (e.g. MKP3 and CA MEK) were fused at the N-terminus of GLIMPSe.
